# Single-cell transcriptomic landscape of cardiac neural crest cell derivatives during embryonic and neonatal development

**DOI:** 10.1101/759118

**Authors:** Xuanyu Liu, Wen Chen, Wenke Li, Ziyi Zeng, James R. Priest, Zhou Zhou

**Author notes:** X.L. and W.C. contributed equally to this manuscript. **Address correspondence to:** Dr. Zhou Zhou, No. 167, Beilishi Road, Xicheng District, Beijing, 100037, People’s Republic of China.

## Abstract

**Rationale:** Cardiac neural crest cells (CNCCs) contribute greatly to cardiovascular development. A thorough understanding of the cell lineages, transcriptomic states and regulatory networks of CNCC derivatives during normal development is essential for deciphering the pathogenesis of CNCC-associated congenital anomalies. However, the transcriptomic landscape of CNCC derivatives during development has not yet been examined at a single-cell resolution.

**Objective:** We sought to systematically characterize the cell lineages, define the developmental chronology and elucidate the transcriptomic dynamics of CNCC derivatives during embryonic and neonatal development.

**Methods and Results:** We performed single-cell transcriptomic sequencing of 34,131 CNCC-derived cells in mouse hearts from eight developmental stages between E10.5 and P7. Through single-cell analyses and single-molecule fluorescence *in situ* hybridization, we confirmed the presence of CNCC-derived mural cells. Furthermore, we found the transition from CNCC-derived pericytes to microvascular smooth muscle cells, and identified the genes that were significantly regulated during this transition through pseudo-temporal analysis. CNCC-derived neurons first appeared at E10.5, which was earlier than previously recognized. In addition, the CNCC derivatives switched from a proliferative to a quiescent state with the progression of development. Gradual loss of the neural crest molecular signature with development was also observed in the CNCC derivatives. Our data suggested that many CNCC-derivatives had already committed or differentiated to a specific lineage when migrating to the heart. Finally, we characterized some previously unknown subpopulations of CNCC derivatives during development. For example, we found that *Penk*+ cells, which were mainly localized in outflow tract cushions, were all derived from CNCCs.

**Conclusions:** Our study provides novel insights into the cell lineages, molecular signatures, developmental chronology and state change dynamics of CNCC derivatives during embryonic and neonatal development. Our dataset constitutes a valuable resource that will facilitate future efforts in exploring the role of CNCC derivatives in development and disease.

## INTRODUCTION

Neural crest cells (NCCs) are a multipotent, migratory cell population that delaminates from the dorsal part of the neural tube via epithelial-to-mesenchymal transition.^1^ During embryogenesis, migratory NCCs give rise to a plethora of cell lineages, and contribute to the development of a variety of tissues and organs, such as the skull bones, adrenal gland, enteric nervous system and heart.^2^ While the heart is mostly of mesodermal origin, NCCs, which are ectodermal derivatives, contribute greatly to heart development.^3^ The subpopulation of NCCs contributing to the heart are referred to as cardiac neural crest cells (CNCCs).^4^ Since these cells were first discovered by Kirby et al.,^5^ CNCCs have been demonstrated to play essential roles in cardiovascular development including the remodeling of the pharyngeal arch arteries, cardiac outflow tract (OFT) septation, valvulogenesis and cardiac innervation.^6^ Genetic or environmental disturbance of the migration, survival and differentiation of CNCCs may result in congenital cardiovascular anomalies. Various human syndromes involving severe congenital heart defects have been associated with CNCCs, such as DiGeorge, Noonan and CHARGE syndromes.^4,6^ A thorough understanding of the cell lineages, transcriptomic states and regulatory networks of CNCC derivatives during normal development is essential for deciphering the pathogenesis of these CNCC-associated congenital cardiovascular anomalies.

In recent decades, significant advances in understanding the CNCC contributions to heart development have been made by using lineage tracing mouse models such as *Wntl-Cre* mice,^7^ although some aspects remain contentious. In these models, all CNCCs and their derivatives are genetically labeled by the *Cre-loxP* recombinase system and observed via LacZ staining or fluorescence imaging (imaging-based lineage tracing).^8^ After delamination from the neural tube (embryonic day 8.5, E8.5), CNCC derivatives first colonize the pharyngeal arch artery and ultimately differentiate into vascular smooth muscle cells (VSMCs) of the aortic arch.^4^ Starting at E10.5, CNCC-derived mesenchymal cells migrate into the OFT and join the cushion mesenchyme, where they participate in the formation of the aorticopulmonary septum for complete separation of the pulmonary and systemic circulation.^9^ These CNCC-derived mesenchymal cells eventually give rise to part of the smooth muscle walls of the great arteries.^10^ The remodeling of OFT cushions also result in the formation of semilunar valvular leaflets, among which CNCC derivatives mainly contribute to the two leaflets adjacent to the aorticopulmonary septum.^7,11^ CNCCs have also been suggested to directly contribute to the smooth muscle walls of the proximal coronary arteries.^7,12^ In addition to the entry point described above (i.e., the arterial pole of the heart), CNCC derivatives enter the heart from a second entry point, the venous pole at E12.5, whereby they penetrate the heart and migrate into the atrioventricular valves.^11,13^ All the melanocytes in atrioventricular valves are derived from CNCCs.^11^

In addition, neurons and glial cells derived from CNCCs contribute to the parasympathetic innervation of the heart.^11,14^ CNCC-derived neurons in the heart were first observed at E11.5.^14,15^ Although it has been suggested that CNCCs are required for normal development of the cardiac conduction system (CCS), it remains contentious whether CNCCs directly contribute to the CCS, which is known to be derived from the myocardium (myocardial conducting cells).^4,6^ Likewise, the presence of CNCC-derived myocardial cells remains controversial, although it has been suggested that CNCCs are essential for normal myocardial development.^4,16^

Mural cells include pericytes that discontinuously ensheath capillaries and microvascular smooth muscle cells (mVSMCs) that cover larger-caliber vessels of the microcirculation as well as their transitional cells.^17,18^ Neural crest-derived mural cells have been identified in various organs, such as the brain, retina, head and thymus.^19,20^ However, one previous study did not find any CNCC-derived mural cells in the heart ventricle at E14.5 using lineage tracing.^21^ Given that only one developmental stage was examined in the previous study, it remains an open question whether CNCCs contribute to cardiac mural cells.

It has become increasingly evident that even at the beginning of migration from the neural tube, the NCCs are heterogeneous, comprising multipotent cells, cells whose differentiation potential are restricted to varying degrees (fate-restricted cells), and even precursors committed to a particular lineage.^22^ Although transcriptomic states have been investigated for pre-migratory or early migrating NCCs in the dorsal neural tube of the chick embryo,^23,24^ little is known regarding the states of CNCC derivatives with respect to their differentiation potential and proliferative ability when they arrive at the heart or during embryonic and neonatal development of the heart.

One obvious drawback of imaging-based lineage tracing is that it cannot provide detailed molecular information about cell state transitions.^8^ Recent technical advances in large-scale single-cell RNA-seq have enabled the transcriptomes of tens of thousands of cells to be assayed at a single-cell resolution.^25^ As a complement to conventional imaging-based lineage tracing, large-scale single-cell RNA-seq allows unbiased cellular heterogeneity dissection, molecular signature identification and developmental trajectory reconstruction at an unprecedented scale and resolution. Large-scale time-series single-cell RNA-seq is becoming a powerful tool for studying the development of complex tissues, organs and even whole organisms.^26,27^ However, to our knowledge, the transcriptomic landscape of CNCC derivatives during embryonic and neonatal development has not yet been examined at a single-cell resolution.

Here, we performed single-cell RNA-seq of CNCC derivatives in mouse hearts from eight developmental stages between E10.5 and P7 (postnatal day 7). We sought to systematically characterize the cell lineages, define the developmental chronology and elucidate the transcriptomic dynamics of CNCC derivatives during embryonic and neonatal development.

## METHODS

The raw sequencing reads have been deposited in the Sequence Read Archive and are available through project accession number PRJNA562135. Detailed methods can be found in the Supplementary Methods.

## RESULTS

### Single-cell transcriptomic sequencing of CNCC derivatives during embryonic and neonatal development

To investigate the transcriptomic landscape of CNCC derivatives during development, we used the *Wnt1-Cre;Rosa26-tdTomato* mouse model to specifically label CNCC-derived cells (Figure 1A). Whole hearts were dissociated, and *tdTomato*-positive cells were sorted for single-cell capture. The developmental stages we selected spanned from the very early time when CNCC derivatives arrived at the cardiac OFT during embryonic development (i.e., E10.5)^9^ to neonatal stage P7 (Figure 1B). 10X Genomics Chromium Single Cell 3’ transcriptomic sequencing libraries were constructed and subjected to sequencing. The sequencing quality metrics were similar across samples, reflecting relatively little technical variation (Online Table I). After the application of stringent quality control, we obtained high-quality single-cell transcriptomes of 34,131 CNCC-derived cells from eight stages. To facilitate further data exploration, we developed a web-based interface for our dataset (http://scrnaseqcncc.fwgenetics.org) that permits interactive examination of expression for any gene of interest.

**Figure 1.**
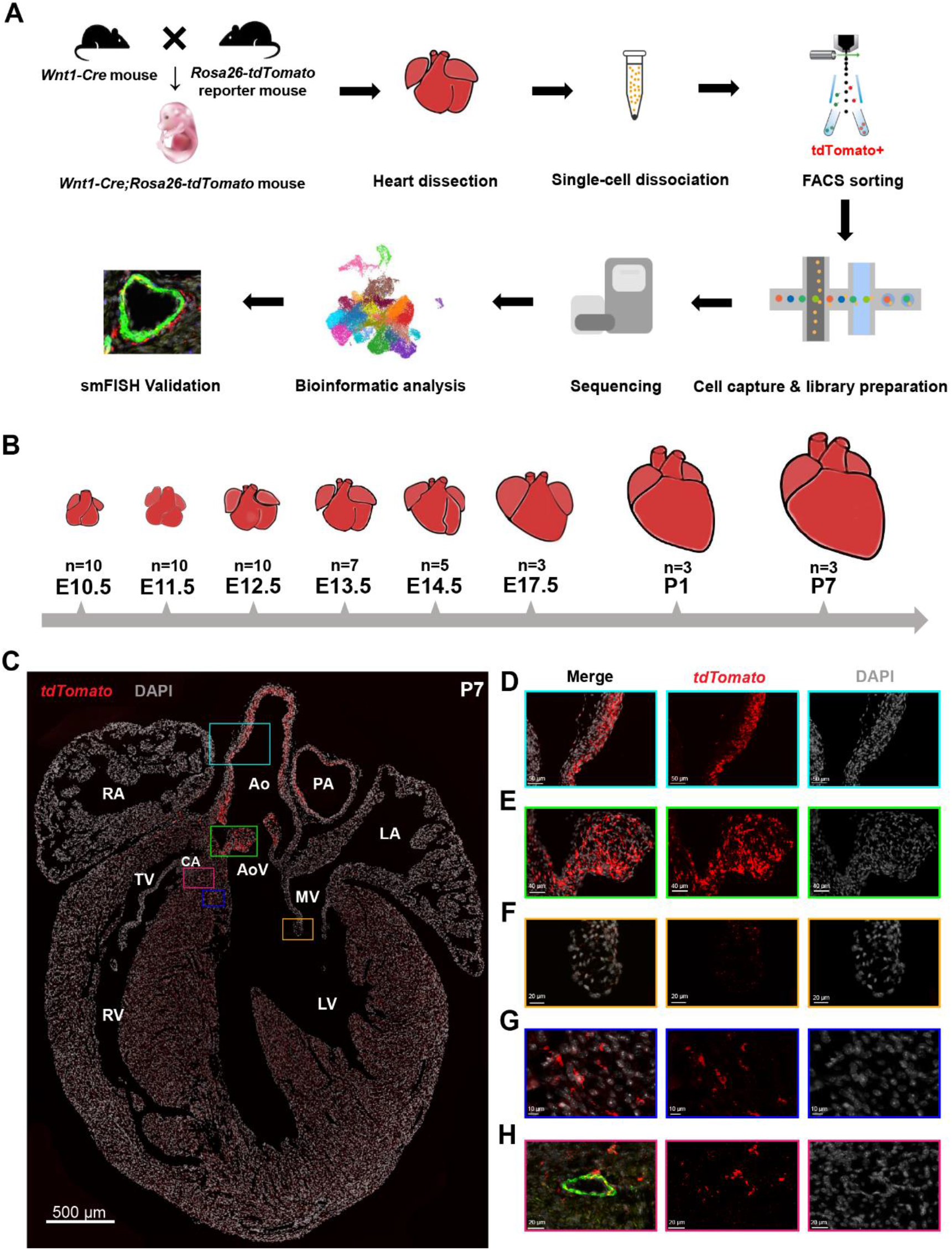
Single-cell RNA-seq and spatial distribution of the CNCC derivatives. **(A)** Schematic representation of the experimental procedure. **(B)** The developmental stages at which the hearts were sampled. Multiple hearts were pooled as a sample for each stage. **(C)** The spatial distribution of the CNCC derivatives labeled by *tdTomato*. **(D-H)** Magnified views of the rectangular regions in C. The same regions are indicated by the same colors. In H, the vessel wall is indicated by the green fluorescence of the VSMC marker *Myh11*. Ao, aorta; AoV, aortic valve; CA, coronary artery; LA, left atrium; LV, left ventricle; MV, mitral valve; PA, pulmonary artery; RA, right atrium; RV, right ventricle; TV, tricuspid valve

### The spatial distribution of the CNCC derivatives

The visualization of *tdTomato*-positive cells by single-molecule fluorescence *in situ* hybridization (smFISH) enabled us to accurately obtain spatial distribution information for the CNCC derivatives, which was not included in the single-cell RNA-seq data (Figure 1C). At neonatal stage P7, we observed a large number of *tdTomato*-positive cells in the walls of the aorta and pulmonary artery (Figure 1C-D) as well as the aortic and pulmonary valve leaflets (Figure 1E, Online Figure I), reflecting the great contribution of CNCCs to the OFT development. Notably, our smFISH results indicated that CNCC-derived cells only populated the inner medial cells of both the ascending aorta and aortic root (Figure 1C-D), thus supporting the view put forth in the most recent report about the distribution of CNCC- and SHF-derived VSMCs.^10^ Consistent with previous reports,^7,11^ the CNCC derivatives were found mainly in the two leaflets adjacent to the aorticopulmonary septum of aortic and pulmonary valves (i.e., right and left leaflets) (Online Figure I). Compared with the aortic and pulmonary valves (Figure 1E), the CNCCs made a much smaller contribution to the atrioventricular valves (Figure 1F). In addition, *tdTomato*-positive cells were found to be embedded in the walls of ventricles (Figure 1G). CNCC-derived VSMCs were observed in the coronary vasculature, as evidenced by the co-expression of *tdTomato* and a specific marker for mature VSMCs (i.e., *Myh11*) (Figure 1H).

### Cell lineages and transcriptomic states of the CNCC derivatives during embryonic and neonatal development

After recognizing the spatial distribution of the CNCC derivatives, we systematically dissected the cell lineages and transcriptomic states of the CNCC derivatives. The unsupervised clustering of the 34,131 CNCC-derived cells from eight stages identified 21 cell clusters (Figure 2A). The data structure was visualized in a two- or three-dimensional UMAP embedding (Online Data I). Six cell lineages were revealed by hierarchical clustering of the clusters based on the average expression of 2,000 selected features (Figure 2B) and the expression of established markers (Figure 2C). The representative molecular signatures for each cluster are shown in Figure 2D (Online Table II).

**Figure 2.**
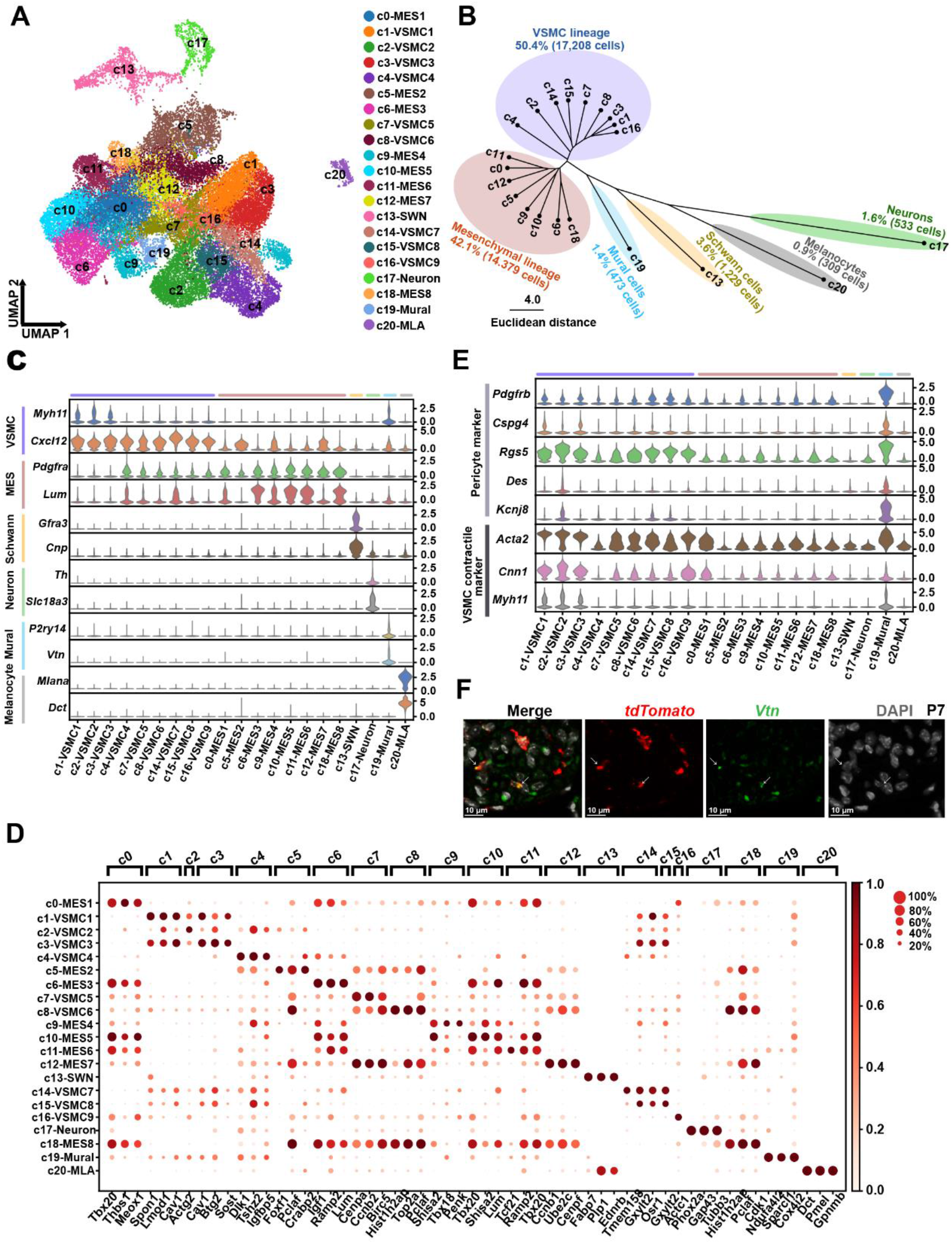
Cell lineages and transcriptomic states of CNCC derivatives during embryonic and neonatal development. **(A)** Single-cell transcriptomes of 34,131 CNCC derivatives projected on a two-dimensional UMAP embedding. Clusters are distinguished by different colors. **(B)** Hierarchical clustering of the clusters based on the average expression of 2,000 selected features. **(C)** Expression of established markers for each cell lineage in each cluster. **(D)** Representative molecular signatures for each cluster. The area of the circles denotes the proportion of cells expressing the gene, and the color intensity reflects the expression intensity. **(E)** Expression of canonical pericyte markers and VSMC contractile markers. **(F)** smFISH validation of CNCC-derived pericytes. Hearts from P7 mice were used. Arrows indicate the co-expression of *tdTomato* and the pericyte marker *Vtn*. MES, mesenchymal cell; MLA, melanocyte; SWN, Schwann cell; UMAP, uniform manifold approximation and projection; VSMC, vascular smooth muscle cell

The VSMC (marked by the mature VSMC marker *Myh11*^28^ and the immature VSMC marker *Cxcl12*^29^) and mesenchymal (marked by *Pdgfra*^30^ and *Lum*^31^) lineages constituted the two largest lineages of the CNCC derivatives (accounting for 50.4% and 42.1% of the derivatives, respectively). Consistent with the differentiation of mesenchymal cells into VSMCs during development, these two lineages were aligned closely in the UMAP embedding (Figure 2A), and some intermediate subpopulations, such as c4, expressed markers of both lineages (Figure 2C). As expected, we identified CNCC-derived neurons (marked by the parasympathetic neuron marker *Slc18a3* and the sympathetic neuron marker *Th*^14^), Schwann cells (marked by *Gfra3* and *Cnp*^32^) and melanocytes (marked by *Mlana*^33^ and *Dct*^34^). We did not find any CCS or myocardial cell clusters, so our data do not support a direct contribution of CNCCs to the CCS and myocardium in the mouse.

Intriguingly, we observed a cluster of mural cells (i.e., c19) based on the pericyte markers recently reported from single-cell studies: *P2ry14*^32^ and *Vtn*^35^ (Figure 2B-C), thus supporting the existence of CNCC-derived mural cells. To further confirm their mural cell identity, we examined the expression of canonical pericyte markers and VSMC contractile markers (Figure 2E). Compared with the others, the c19 cluster expressed higher levels of canonical pericyte markers including *Pdgfrb*, *Cspg4* (NG2), *Rgs5*, *Des* and *Kcnj8*.^18,36^ It also exhibited high expression of VSMC contractile markers, including *Acta2*, *Cnn1* and *Myh11*,^28^ reflecting a heterozygous microvascular mural population comprising both pericytes and mVSMCs. Our smFISH results ultimately validated the presence of CNCC-derived pericytes in the heart through the co-expression of *tdTomato* and *Vtn* (Figure 2F).

### CNCC-derived pericytes transition to microvascular smooth muscle cells

To understand the heterogeneity of the CNCC-derived mural cells, we performed subclustering of the c19 cells from stage P7 (227 cells, accounting for 48% of the cluster). Two subclusters were identified, sc1 and sc2, which correspond to pericytes and mVSMCs, respectively, based on the expression of markers (Figure 3A-B). RNA velocity analysis represents a computational framework that can infer the direction and rate of cellular state changes based on the relative abundance of spliced and unspliced transcripts.^37^ Our RNA velocity analysis revealed a transition from pericytes to mVSMCs, which is in agreement with a previous report that pericytes serve as progenitors for smooth muscle cells of the coronary vasculature.^38^

**Figure 3.**
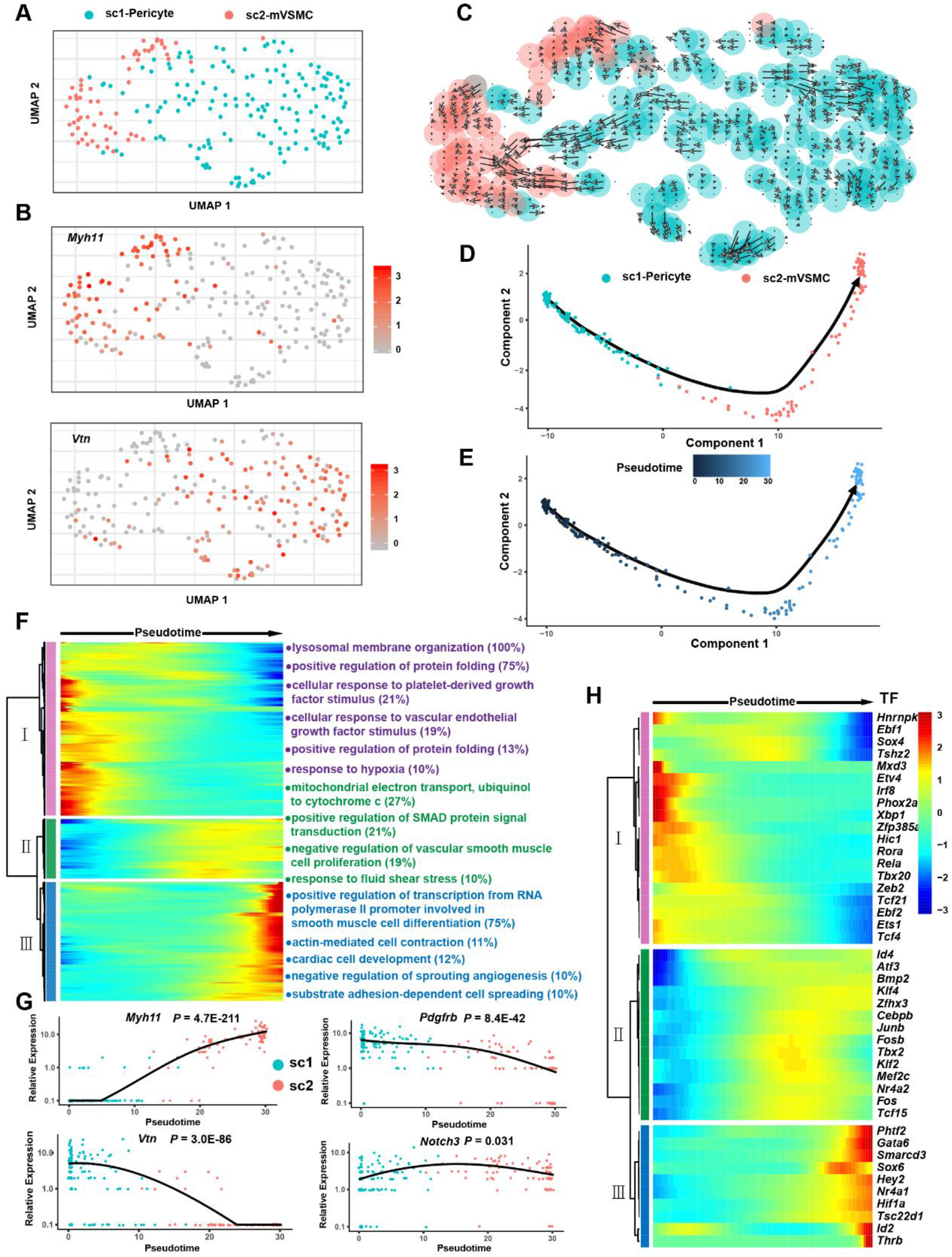
Cellular transition from CNCC-derived pericytes to microvascular smooth muscle cells. **(A)** Subclustering of the mural cell cluster c19 reveals two subpopulations. mVSMC, microvascular smooth muscle cells. Only P7 cells were considered in this analysis. **(B)** The expression distribution of the VSMC-specific marker *Myh11* and the pericyte marker *Vtn*. **(C)** RNA velocity analysis reveals a transition from CNCC-derived pericytes to mVSMCs. The direction and length of the arrow reflect the direction and rate of cellular state changes, respectively. **(D)** Linear trajectory constructed via pseudo-temporal ordering of cells. **(E)** Transition trajectory colored according to pseudotime. **(F)** Hierarchical clustering of the genes that were significantly regulated during the progression of the transition. Only genes with an adjusted P-value <1E-04 are shown here. The number in the parentheses represents the percentage of genes associated with the Gene Ontology term for which the gene cluster is significantly enriched (adjusted P-value <0.05). **(G)** The expression changes in *Myh11, Vtn, Pdgfrb* and *Notch3* during the progression of the cellular transition. **(H)** Transcriptional factors that were significantly regulated during the transition.

The single-cell data provided a unique opportunity for interrogating the regulatory changes during the transition. Pseudo-temporal ordering of the cells using Monocle2 resulted in the construction of a linear trajectory of cellular transition (Figure 3D-E). We further identified 952 genes that were significantly regulated during the progression of the transition (Figure 3F, Online Table III, adjusted P-value <1E-04). Hierarchical clustering of the identified genes revealed three clusters. Gene cluster I represented the molecular characteristics of pericytes and was mainly enriched for lysosomal membrane organization, cellular response to platelet-derived growth, cellular response to vascular endothelial growth factor stimulus and response to hypoxia (Figure 3F, Online Table IV). Gene cluster II reflected the phenotype of transitioning cells and was mainly enriched for mitochondrial electron transport, positive regulation of SMAD protein signal transduction, negative regulation of vascular smooth muscle cell proliferation and the response to fluid shear stress. Gene cluster III represented the characteristics of mVSMCs and was mainly enriched for positive regulation of transcription from RNA polymerase II promoter involved in smooth muscle cell differentiation and substrate adhesion-dependent cell spreading. Figure 3G shows the expression dynamics of the pericyte markers *Vtn* and *Pdgfrb* as well as the VSMC marker *Myh11*, reflecting a continuum of phenotypic changes in cells embedded in the walls of the microvasculature. Interestingly, Notch3 signaling has been suggested to be important in the pericyte to VSMC transition.^38^ We found that *Notch3* was significantly up-regulated specifically in the middle phase of the trajectory (Figure 3G, adjusted P-value = 0.031), while other Notch receptors including *Notch1, Notch2* and *Notch4* were not significantly regulated. Moreover, we examined the transcription factors (TFs) that were significantly regulated during the transition (Figure 3H). Notably, *Fosb*, *Tbx2* and *Klf2* were specifically up-regulated in the middle phase of the trajectory, implying that they played roles in the transition.

### Developmental chronology and transcriptomic state change dynamics of CNCC derivatives during development

The study of the developmental chronology of CNCC derivatives has previously been limited by improper or limited cell markers for each developmental stage. The large-scale single-cell RNA-seq dataset gave us an unpreceded opportunity, since the cells were clustered in an unbiased manner based on the whole transcriptome, without the need for *a priori* knowledge about the cell markers. Figure 4A and 4B show the proportion of each cluster in each stage and the proportion of cells from each stage in each cluster, respectively. As shown in cluster I of Figure 4B, the melanocyte lineage first appeared at E11.5 and then greatly expanded at E14.5. These results are consistent with a previous report that *Dct* (a melanocyte marker) expression is first observed at E11.5 and that a larger number of melanocytes are found in the atrioventricular endocardial cushions at E14.5,^34^ reflecting the reliability of our dataset. Surprisingly, CNCC-derived neurons were found to appear first at E10.5, which was earlier than previously recognized (E11.5).^13,14^ Only six E10.5 neuron cells were captured, but they all expressed neural markers (Online Table V), thus excluding errors from data integration and clustering.

**Figure 4.**
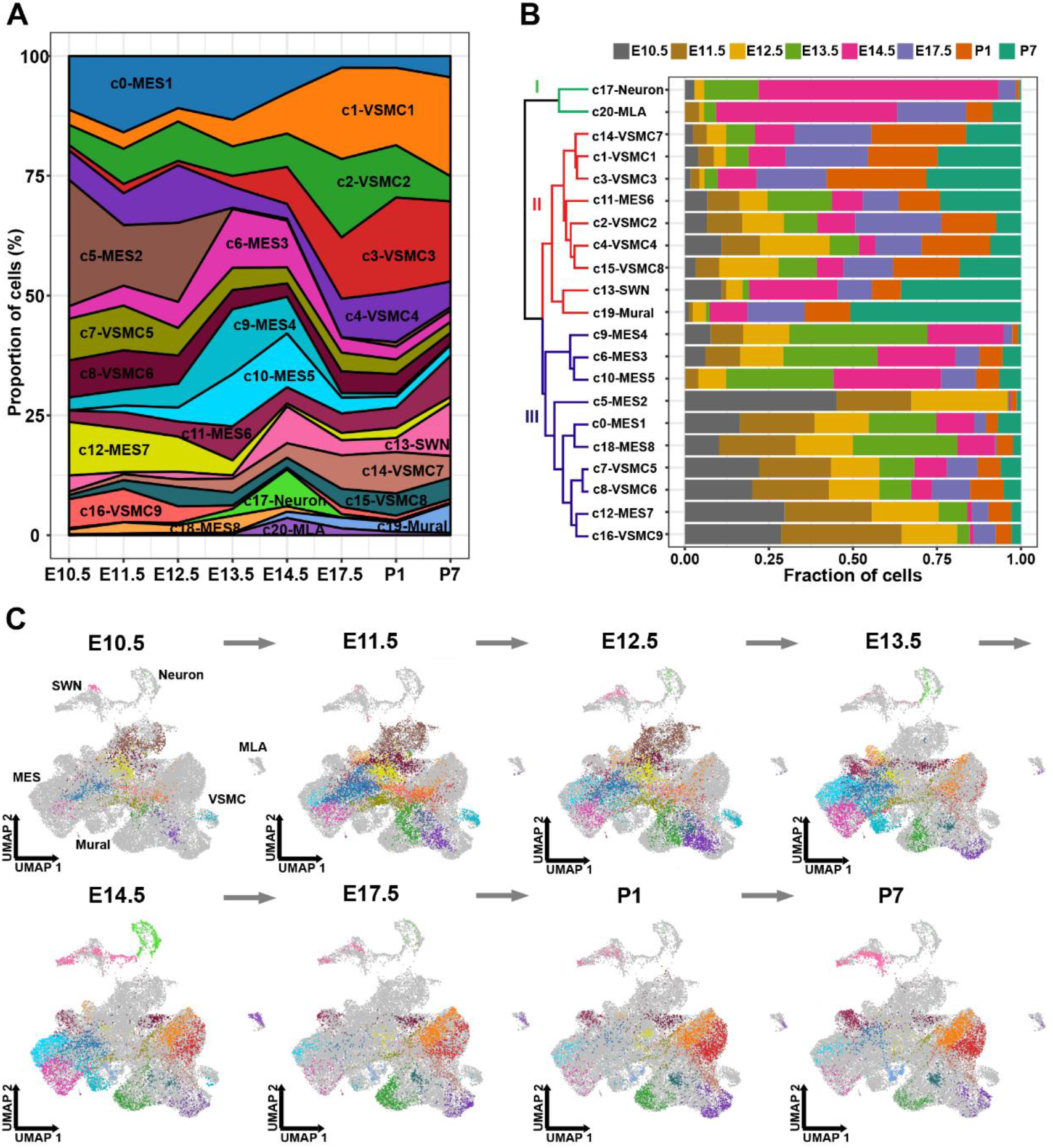
The developmental chronology and transcriptomic state change dynamics of CNCC derivatives. **(A)** The proportion of cells of each cluster in each stage. **(B)** The proportion of cells from each stage in each cluster. All samples are normalized to the same number of cells (2,026). The dendrogram shows the hierarchical clustering of the cell clusters based on the proportion of cells from each stage. **(C)** Dynamic changes in the transcriptomic states of CNCC derivatives during development. The cells are colored according to the clusters. MES, mesenchymal cell; MLA, melanocyte; SWN, Schwann cell; UMAP, uniform manifold approximation and projection; VSMC, vascular smooth muscle cell

As expected, the VSMC lineage expanded mainly at the later stages of development (after E14.5; cluster II in Figure 4B), and the mesenchymal lineage expanded mainly at the early stages (before E14.5; cluster III in Figure 4B). Notably, the mural cells expanded greatly postnatally (especially at P7), in line with the increase in capillary growth during the postnatal development of the heart.^39^ The transcriptomic state change dynamics during development of the CNCC-derived lineages can be clearly visualized in Figure 4C. Notably, the earliest sample from E10.5 included multiple cell lineages, supporting the view that many CNCC-derivatives had already committed or differentiated to a specific lineage when they arrived at the heart.

### Gradual loss of proliferation and the neural crest molecular signature with development in CNCC derivatives

We further characterized the CNCC derivatives with respect to their proliferation ability and differentiation potential when they arrived at the heart as well as during embryonic and postnatal development. We found that the CNCC derivatives were highly proliferative when they arrived at the heart (E10.5) and switched from a proliferative to a quiescent state with the progression of development (Figure 5A). Some clusters, such as c18, c12, c8, c5 and c7, were highly proliferative (Figure 5B). We further investigated the differentiation potential of CNCC derivatives by examining the expression of a list of markers for pluripotency and pre-migratory neural crest cells that was compiled by a previous study.^23^ No cell clusters were found to exhibit high expression of pluripotency genes such as *Nanog* and *Pou5f1* (*Oct4*), suggesting that the CNCC-derived cells generally did not possess stemness after migrating into the heart (Figure 5C). Moreover, we observed gradual loss of the neural crest molecular signature with development in the CNCC derivatives (Figure 5D). Notably, the CNCC-derived cell lineages exhibited differences in the neural crest molecular signature (Figure 5E). Surprisingly, the melanocytes, rather than the mesenchymal cells, were most similar to pre-migratory neural crest cells. The melanocytes also showed high expression of other neural crest markers, including *Pax3* and *Kit*^40^ (Online Figure II).

**Figure 5.**
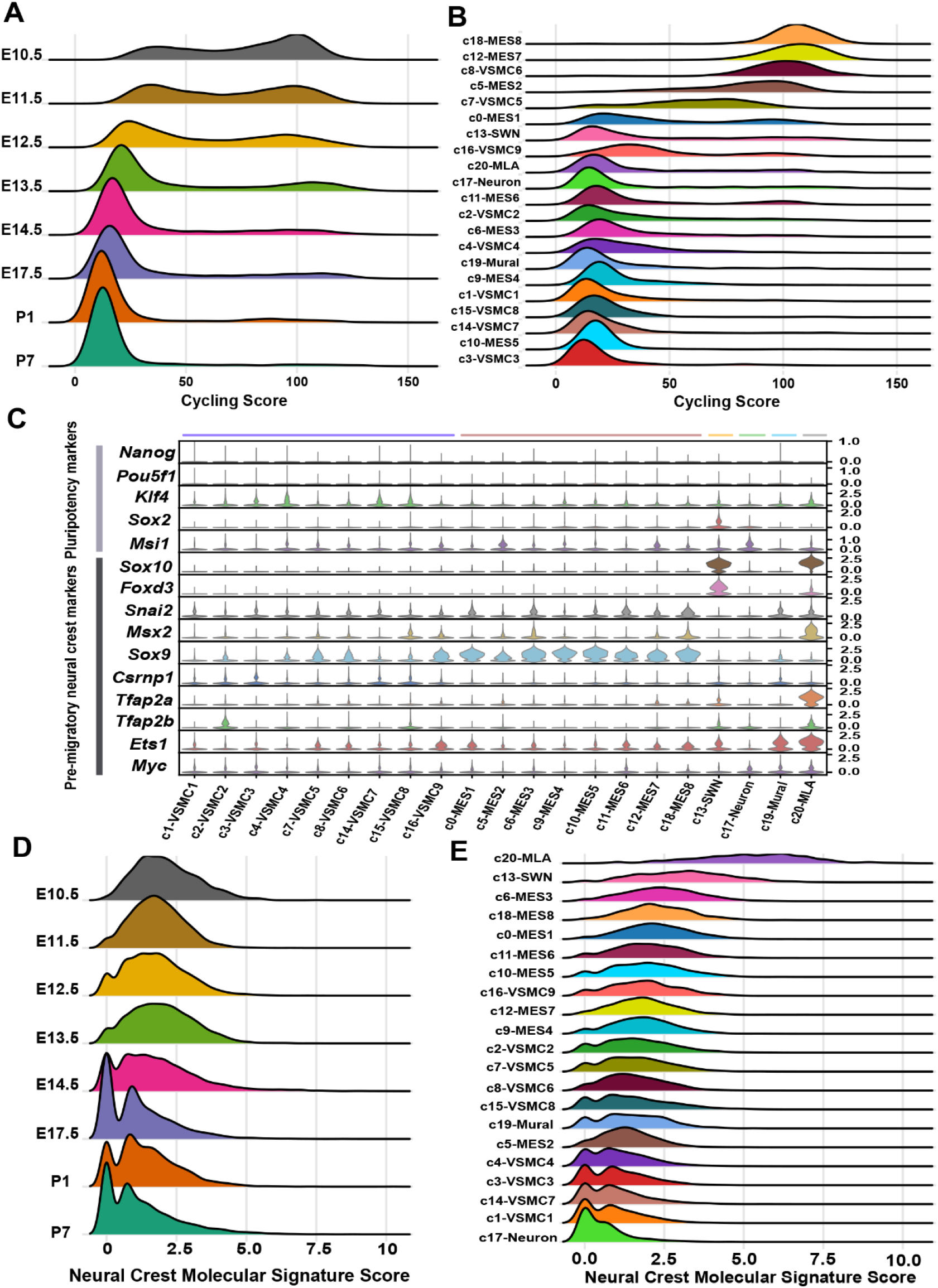
Gradually loss of proliferation and the neural crest molecular signature with development in the CNCC derivatives. **(A)** Ridge plot showing that CNCC derivatives switched from a proliferative to a quiescent state with the progression of development. The cycling score of each cell was calculated by summing the log-normalized expression of the cycling genes (“g2m.genes” and “s.genes” in Seurat). **(B)** Ridge plot showing the proliferative ability of each cell cluster. **(C)** The expression of pluripotency and pre-migratory neural crest markers in each cell cluster. **(D)** Ridge plot showing that the divergence in molecular signatures between the CNCC derivatives and pre-migratory neural crest cells increased during development. The neural crest molecular signature score was calculated by summing the log-normalized expression of the pre-migratory neural crest markers (shown in C). **(E)** Ridge plot showing the distribution of the neural crest molecular signature score in each cluster.

### Characterization of interesting cell subpopulations of CNCC derivatives during development

Single-cell RNA-seq permits the identification of previously unrecognized subpopulations, and we characterized some interesting subpopulations that deserve further study. Cluster c1, c2 and c3 exhibited high expression of *Myh11*, thus representing relatively mature VSMCs (Figure 2C). However, c2 was aligned distant from c1 and c3 in the UMAP space (Figure 2A, Online Data I), suggesting that c2 represents another branch of the VSMC lineage distinct from c1 and c3, while the last two clusters aligned together closely. Interestingly, compared with c1 and c3, c2 expressed significantly higher levels of contractile markers such as *Myh11* and *Cnn1* as well as pericyte markers such as *Rgs5* and *Kcnj8* (Figure 6A). Cluster c1 and c3 expressed significantly higher levels of extracellular matrix genes such as *Eln* and *Fbln2* than c2. Taken together, c2 may represent the CNCC-derived VSMCs of the coronary vasculature, while c1 and c3 may represent the CNCC-derived VSMCs of the great arteries. The smFISH results confirmed that the coronary arteries expressed significantly higher levels of *Myh11* than the great arteries (Figure 6B-C).

**Figure 6.**
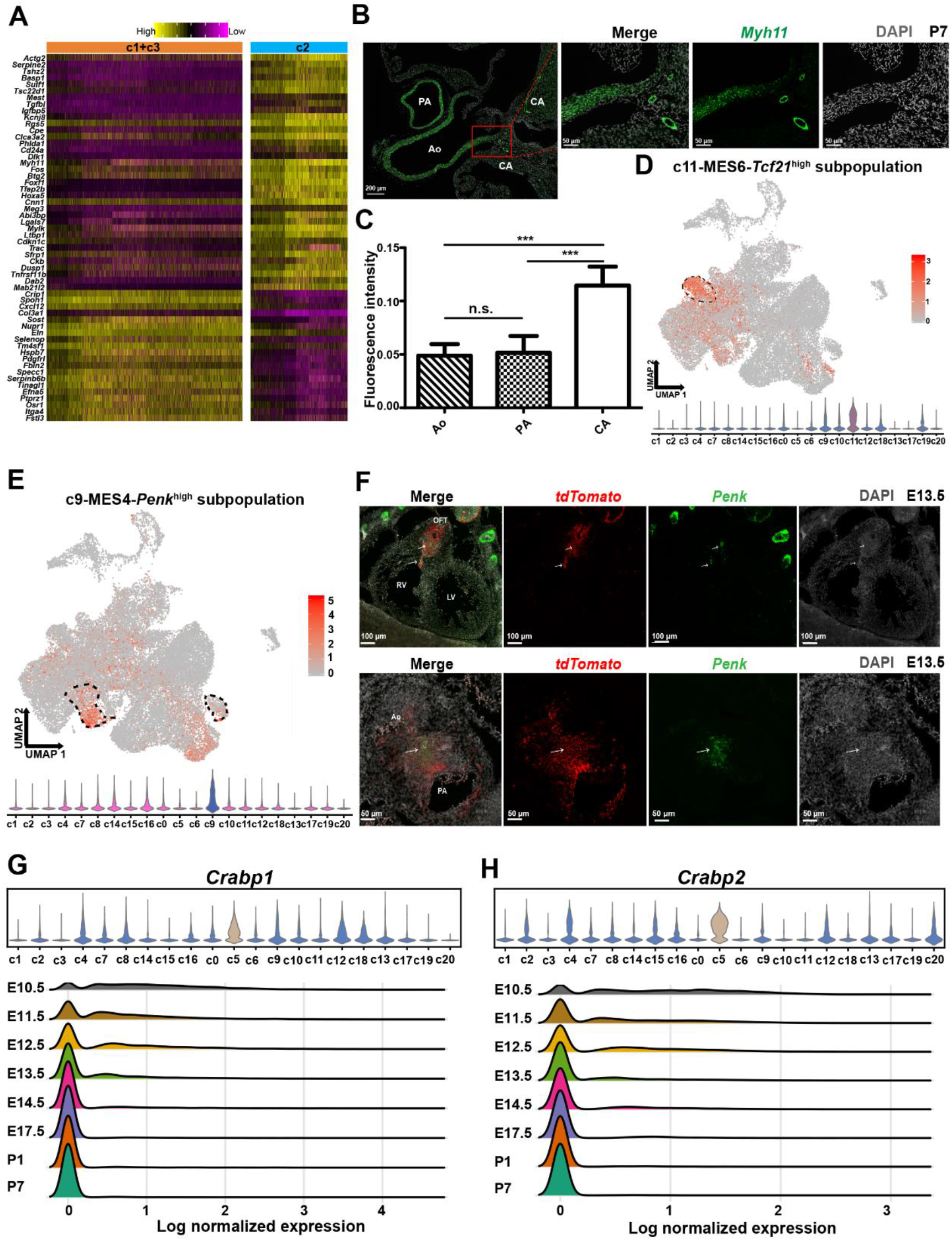
Characterization of interesting cell subpopulations of CNCC derivatives during development. **(A)** Heatmap showing the difference between VSMC clusters c2 and c1+c3. The significance threshold was set to an adjusted P-value < 0.05 and log2-fold change > 0.25. **(B)** smFISH results showing significantly higher expression of the contractile marker *Myh11* in VSMCs of the coronary arteries than in VSMCs of the great arteries. **(C)** Quantitative analysis of the fluorescent intensity of *Myh11* expression confirms significantly higher expression of *Myh11* in VSMCs of the coronary arteries than in VSMCs of the great arteries. The bar height represents the average intensity of five biological replicates (±SE). One-way ANOVA with Turkey’s post-hoc test. P-value≤ 0.001 (***). n.s. not significant. **(D)** Mesenchymal cluster c11 with high *Tcf21* expression may represent valve interstitial cells. **(E)** Mesenchymal cluster c9 shows high expression of *Penk*. **(F)** smFISH results showing that the *Penk*+ cells are mainly localized in the OFT region and are derived from CNCC. **(G)** *Crabp1* shows high expression in mesenchymal cluster c5 and its expression decreases with development. **(H)** *Crabp2* shows high expression in mesenchymal cluster c5 and its expression decreases with development. Ao, aorta; CA coronary artery; LV, left ventricle; OFT, outflow tract; PA, pulmonary artery; RV, right ventricle.

The mesenchymal cluster c11 showed high expression of the transcription factor *Tcf21* (Figure 6D), which may indicate a subpopulation of CNCC-derived valve interstitial cells based on the previous reports.^33,41^ Our lab previously identified a *Penk*^+^ mesenchymal subpopulation in the developing OFT; however, whether this subpopulation is derived from CNCCs was not answered.^29^ In this study, we found the mesenchymal cluster c9 showed high expression of *Penk* (Figure 6E), and the smFISH results showed that the *Penk*^+^ cells were mainly localized in the OFT cushions where the aortopulmonary septum formed, and all the *Penk+* cells were derived from CNCCs (Figure 6F).

Another interesting mesenchymal cluster is c5, which mainly comprised cells from the early stages (Figure 3A), suggesting that it may represent the early state of the CNCC-derived mesenchymal cells. This subpopulation showed high expression of the *Crabp1* gene, encoding cellular retinoic acid binding protein 1 (Figure 6G), which has been reported as the top marker of CNCC-derived mesenchymal cells at E9.25.^42^ It also showed high expression of the *Crabp2* gene, encoding cellular retinoic acid binding protein 2 (Figure 6H). The expression of *Crabp1* and *Crabp2*, two important regulators of retinoic acid signaling, decreased during development. These results indicate that CNCC derivatives are more sensitive to retinoic acid signaling at early stages of development.

## DISCUSSION

The neural crest is fascinating. The formation, migration and differentiation of NCCs and NCC-associated pathologies have been the subject of intense research since the discovery of these cells 150 years ago.^43^ CNCCs play critical roles in the evolution and development of the vertebrate cardiovascular system.^44^ In this study, we systematically investigated the transcriptional landscape of CNCC derivatives during cardiac development at a single-cell resolution. On the basis of large-scale single-cell RNA-seq analyses and smFISH validation, we reported the presence of CNCC-derived mural cells associated with the microvasculature. Furthermore, we found the transition from CNCC-derived pericytes to mVSMCs and identified the genes that were significantly regulated during the transition through pseudo-temporal ordering analysis. We defined the developmental chronology of the CNCC-derived lineages and found that the CNCC-derived neurons first appeared at E10.5, which was earlier than previously recognized. Our data indicated that many CNCC derivatives had already committed or differentiated to a specific lineage when they arrived at the heart. We found that the CNCC derivatives were highly proliferative when migrating into the heart, and switched from a proliferative to a quiescent state with the progression of development. Gradual loss of the neural crest molecular signature with development was also observed in the CNCC derivatives. The CNCC-derived cell lineages exhibited differences in the neural crest molecular signature. Surprisingly, the melanocytes were most similar to the pre-migratory neural crest cells. Finally, we confirmed some interesting subpopulations of the CNCC derivatives during development. For example, we found that *Penk*+ cells were mainly localized in the OFT cushions where the aortopulmonary septum formed, and confirmed that all the *Penk*+ cells were derived from CNCCs.

Understanding the origin and regulators driving the development of the cardiac vasculature is an important topic in developmental biology. Microvascular mural cells, comprising microvascular pericytes and microvascular smooth muscle cells, have recently been recognized playing a critical role in cardiac vascular homeostasis and disease.^18^ The plasticity of microvascular pericytes makes them promising cells for application in cardiac regenerative medicine.^17^ Nevertheless, the phenotypes of microvascular mural cells are variable, and canonical markers such as *Pdgfrb, Des* and *Cspg4* do not specifically label them, as also shown in our data (Figure 2E), thus impeding the identification of the source and role of this important but heterogeneous population of cells. Using lineage tracing and canonical markers, previous studies have reported the embryonic origin of cardiac mural cells from epicardium or endocardial endothelial cells.^21,45^ Based on single-cell clustering and novel markers recently reported from single-cell studies, we identified a third source of mural cells in the heart (i.e., CNCC-derived mural cells) (Figure 2B-C). This finding makes sense because NCCs have already been reported to give rise to mural cells in many organs, such as the brain, retina and thymus.^19,20^ We also found that the mural cells expanded greatly postnatally (especially at P7, Figure 4B), in line with the increase in capillary growth during the postnatal development of the heart.^39^ This may be one of the reasons why the CNCC-derived mural cells were not identified in a previous study^21^ since the CNCC-derived mural cells are relatively few at E14.5 (the stage that study only examined). Consistent with the phenotypic heterogeneity of cardiac mural cells, our results reflected a more complex embryonic origin of cardiac mural cells than previously recognized. Whether cardiac mural cells of different origins behave differentially during pathological processes of the coronary vasculature deserves further study. Although a previous study indicated that pericytes can transition to mVSMCs,^38^ the gene expression dynamics underlying this transition are not yet fully elucidated. Through pseudo-temporal ordering of single cells, we confirmed the linear trajectory of the pericyte-to-mVSMC transition and, for the first time, elucidated the previously unknown regulatory changes during the transition (Figure 3F). Our results support the role of Notch3 signaling during the transition and provide candidate regulators potentially driving the process (Figure 3H).

Although our data do not support a direct contribution of CNCCs to the CCS and myocardium, our results highlight the contribution of CNCCs to cardiac vessels of different calibers, from the VSMCs of the great arteries to mural cells wrapping the microvasculature. Moreover, we found that the phenotypes of the cells wrapping the cardiac vessels may vary as a function of the caliber of the vessels. For example, the coronary arteries expressed significantly higher levels of the contractile marker *Myh11* than the great arteries (Figure 6A-C). Our results reflect a continuum of cell phenotypes along the cardiac vascular tree with VSMCs and pericytes at the two ends of the phenotypic spectrum. The heterogeneity of the phenotypes of vessel-associated cells in the brain vasculature has been dissected using single-cell RNA-seq.^36^ The phenotypic heterogeneity of the cardiac vasculature is also complex and deserves to be explored at a single-cell resolution in the future by integrating single-cell RNA-seq data with spatial transcriptomic data.^46^

Due to the limitations of imaging-based lineage tracing used in the previous studies,^7,11^ we know little about the states of CNCC derivatives when they migrate to the heart or the molecular change dynamics during development. The large-scale single-cell RNA-seq dataset gave us unpreceded opportunity to explore these questions. Unexpectedly, CNCC-derived neurons expressing mature neuron markers were found to first appear at E10.5 (Figure 4B, Online Table V), which is earlier than previously recognized.^13,14^ Notably, the earliest sample investigated from E10.5 contained multiple cell lineages and exhibited the expression of lineage-specific mature markers (Figure 4C). The neuron, Schwann and melanocyte lineages aligned relatively distant from the mesenchymal lineage in the UMAP embedding (Figure 4C), suggesting that most cells of these lineages were not differentiated from the mesenchymal lineage after migrating into the heart. No cell clusters were found to highly express the pluripotency genes such as *Nanog* and *Pou5f1*, suggesting that CNCC-derived cells generally do not possess stemness after migrating into the heart (Figure 5C). Taken together, our results support the view that many CNCC-derivatives have already committed or differentiated to a specific lineage when they arrived at the heart. In addition, we observed that the CNCC derivatives were highly proliferative when migrating into the heart, and switched from a proliferative to a quiescent state with the progression of development (Figure 5A). We also observed gradual loss of the neural crest molecular signature with development in the CNCC derivatives. These findings revealed by single-cell analyses provide a deeper understanding of the CNCC derivatives during development.

In conclusion, our study provides novel insights into the cell lineages, molecular signatures, developmental chronology and state change dynamics of CNCC derivatives during embryonic and neonatal development. Our dataset constitutes a valuable resource that will facilitate future efforts to explore the roles of CNCC derivatives in development and disease.

## Supporting information

Online Table III

## AUTHOR CONTRIBUTIONS

X.L. performed data analysis, interpreted the results, and wrote the manuscript. W.C. performed the wet lab experiments with the assistance of Z. Zeng. W.L. designed the web interfaces. J. R. P. gave suggestions on result interpretation. Z. Zhou conceived the project. X.L. and W.C. participated in designing the project.

## DISCLOSURES

There are no conflicts of interest to declare by any of the authors.

## ACKNOWLEDGMENTS

We thank Dr. Zhen Zhang at Children’s Hospital of Shanghai for transferring the Wnt1-Cre mouse line to our lab.

## SOURCES OF FUNDING

This work is supported by grants from the National Natural Science Foundation of China (81900282), the CAMS Initiative for Innovative Medicine (2016-I2M-1-016), the Post-doctoral International Exchange Project (2018-BSH04), and the Foundation for Fuwai Hospital Youth Scholars (2019-F08).

## Nonstandard Abbreviations and Acronyms

CCS: Cardiac conduction system
CNCC: Cardiac neural crest cell
GEM: Gel Beads-in-Emulsion
mVSMC: Microvascular smooth muscle cell
NCC: Neural crest cell
OFT: Outflow tract
PCA: Principal component analysis
smFISH: Single-molecule fluorescence in situ hybridization
TF: Transcriptional factor
UMAP: Uniform manifold approximation and projection
UMI: Unique molecular identifier
VSMC: Vascular smooth muscle cell

## Notes

http://scrnaseqcncc.fwgenetics.org

