## SUPPLEMENTAL METHODS AND FIGURES for "Single-cell transcriptomic landscape of cardiac neural crest cell derivatives during embryonic and neonatal development"

### SUPPLEMENTAL MATERIALS

#### SUPPLEMENTAL METHODS:

##### Mice:

The *Wnt1-Cre;Rosa26-tdTomato* mice were obtained by crossing a *Wnt1-Cre* transgenic line (Jax Labs; 003829) with a *ROSA26-tdTomato* reporter line (Jax Labs; 007909). The *Wnt1-Cre;Rosa26-tdTomato* mice were confirmed by the presence of fluorescence in the heart using a Discovery V20 stereomicroscope equipped with a 550 nm laser and a Texas Red filter set (Carl Zeiss AG). The *ROSA26-tdTomato* reporter line was purchased from Beijing Vitalstar Biotechnology Co., Ltd. The *Wnt1-Cre* mouse line was transferred from Dr. Zhen Zhang's lab at Children's Hospital of Shanghai.

##### Single-cell suspension preparation and cell sorting:

The whole hearts of embryonic or postnatal *Wnt1-Cre;Rosa26-tdTomato* mice were microdissected under a Stemi 305 compact stereo microscope (Carl Zeiss AG, Germany). The dissected hearts were rinsed with cold Dulbecco’s phosphate-buffered saline to remove most of the red blood cells. To ensure a sufficient number of *tdTomato*-positive cells for single-cell capture, we pooled multiple hearts as a sample for each developmental stage (the numbers of hearts are indicated in Figure 1B). Then, the heart tissues were dissociated using a Pierce Primary Cardiomyocyte Isolation Kit (Thermo Fisher Scientific, #88281) according to the manufacturer’s instructions. The obtained single-cell suspensions were filtered using a 40-mm strainer to remove any cell debris or large clumps. The *tdTomato*-positive cells in the whole heart suspensions were sorted using a BD FACSAria™ II cell sorter (BD Biosciences, USA). Thereafter, the cell suspensions were pelleted and washed twice at 400 g for 5 min at 4 °C, and the pellets were resuspended in Hank’s balanced salt solution with 0.04% bovine serum albumin. Cell viability and concentrations were measured with a TC20 automated cell counter (Bio-Rad, USA).

##### Single-cell RNA-seq library preparation and sequencing:

Single-cell Gel Beads-in-Emulsion (GEM) generation, barcoding, post GEM-RT cleanup, cDNA amplification and cDNA library construction were performed using Chromium Single Cell 3′ Reagent Kit v2 chemistry (10X Genomics, USA) following the manufacturer’s protocol. The resulting libraries were sequenced on a NovaSeq 6000 System (Illumina, USA).

##### Sample demultiplexing, barcode processing and UMI counting:

The official software Cell Ranger v3.0.2 (https://support.10xgenomics.com) was applied for sample demultiplexing, barcode processing and unique molecular identifier

(UMI) counting. Briefly, the raw base call files generated by the sequencers were demultiplexed into reads in FASTQ format using the ‘‘cellranger mkfastq’’ pipeline. Then, the reads were processed using the ‘‘cellranger count’’ pipeline to generate a gene-barcode matrix for each library. During this step, the reads were aligned to the mouse (*Mus musculus*) reference genome (version: mm10) and the *tdTomato* sequence. The resulting gene-cell UMI count matrices of all samples were ultimately concatenated into one matrix using the ‘‘cellranger aggr’’ pipeline.

Data cleaning, normalization, feature selection, integration and scaling:

The concatenated gene-cell barcode matrix was imported into Seurat v3.0.2,[1,2] for data preprocessing. To exclude genes likely detected from random noise, we filtered out genes with counts in fewer than 3 cells. To exclude poor-quality cells that might have resulted from doublets or other technical noise, we filtered cell outliers (> third quartile + 1.5 × interquartile range or < first quartile - 1.5 × interquartile range) based on the number of expressed genes, the sum of UMI counts and the proportion of mitochondrial genes. To further remove doublets, we filtered out cells based on the predictions by Scrublet.[3] In addition, cells enriched in hemoglobin gene expression were considered red blood cells and were excluded from further analyses. The sum of the UMI counts for each cell was normalized to 10,000 and log-transformed. For each sample, 2,000 features (genes) were selected using the “FindVariableFeatures” function of Seurat under the default settings. Genes on the sex chromosomes were removed from the list of selected features. To correct for potential batch effects and identify shared cell states across datasets, we integrated all the datasets via canonical correlation analysis (CCA) implemented in Seurat. To mitigate the effects of uninteresting sources of variation (e.g., the cell cycle), we regressed out the mitochondrial gene proportion, UMI count, S phase score and G2M phase score (calculated by the “CellCycleScoring” function) with linear models using the “ScaleData” function. Then, the data were centered for each gene by subtracting the average expression of that gene across all cells, and were scaled by dividing the centered expression by the standard deviation.

Dimensional reduction and clustering:

The integrated data were imported into the Scanpy toolkit[4] for dimensional reduction and clustering. Briefly, the expression of the selected genes was subjected to linear dimensional reduction through principal component analysis (PCA). Then, the first 30 principal components of the PCA were used to compute a neighborhood graph of the cells. The neighborhood graph was ultimately embedded in two- or three-dimensional space using the non-linear dimensional reduction method of uniform manifold approximation and projection (UMAP).[5] The neighborhood graph of cells was clustered using Louvain clustering (resolution=1).[6]

Differential expression and function enrichment analysis:

The gene signature of each cell cluster was obtained using the “scanpy.tl.rank_genes_groups” tool implemented in Scanpy. The significance threshold was set to a corrected P-value < 0.05. The top 100 genes were retained as the gene signature of each cluster. Differentially expressed genes between two groups of cells were detected with the likelihood-ratio test (test.use: "bimod") implemented in the "FindMarkers" function of Seurat. The significance threshold was set to an adjusted P-value < 0.05 and a log2-fold change > 0.25. Functional enrichment analyses of a list of genes were performed using ClueGO[7] with an adjusted P-value threshold of 0.05 and a percentage of genes associated with the Gene Ontology term ≥ 10%.

RNA velocity analysis:

RNA velocity analysis was performed using scVelo v0.1.19 (https://github.com/theislab/scvelo). First, the expression of the spliced and unspliced mRNAs of each gene in each cell was determined separately for each sample using the velocyto v0.17 tool (https://github.com/velocyto-team/velocyto.py). The first 30 PCA components were used to calculate the first- and second-order moments. Then, the gene-specific velocities were estimated using the "scvelo.tl.velocity" function, and velocity graphs were constructed using the "scvelo.tl.velocity_graph" function. Finally, the velocities were projected onto a two-dimensional UMAP embedding.

Pseudo-temporal ordering of single cells along the differentiation trajectory:

Pseudo-temporal ordering of the cells along the differentiation trajectory was performed using Monocle2.[8] Briefly, the ordering was based on 1,000 genes that differed in expression between clusters selected via an unsupervised procedure: "dpFeature". Then, the data space was reduced to two dimensions with the method "DDRTree". The cells were ultimately ordered in pseudotime, and cells exhibiting high expression of pericyte markers were considered to represent the beginning of the trajectory. Once the pseudotime was assigned for each cell, we identified genes that were significantly regulated as differentiation progressed using the "differentialGeneTest" function. The statistically significant threshold was set to a q-value < 1E-04.

Single-molecule fluorescence *in situ* hybridization and quantitative analyses of the fluorescent signal intensity:

To detect the expression of the target gene in vivo, we performed single-molecule fluorescence *in situ* hybridization (smFISH) using RNAscope® technology for formalin-fixed paraffin-embedded (FFPE) tissues. In brief, the dissected embryo or heart tissues were subjected to formalin fixation, alcohol dehydration and paraffin embedding. Then, we cut the embedded tissues into sections of 4 μm in thickness and performed smFISH using the RNAscope® Multiplex Fluorescent Reagent Kit v2 (Cat. No. 323100, Advanced Cell Diagnostics, lnc., USA) according to the manufacturer's instructions. Fluorescent signals were scanned with the Vectra® Polaris™ pathology imaging system (PerkinElmer, USA). The fluorescent signal intensities in different parts of the heart were quantified with the ImageJ v1.48 tool.[9] The probes used for smFISH were as follows: Mm-Myh11 (Cat# 316101), Mm-Pdgfrb (Cat# 411381), Mm-Vtn (Cat# 443601), Mm-Penk (Cat# 318761) and tdTomato (Cat# 317041), along with a negative control probe (Cat# 321831) and a positive control probe (Cat# 321811).

Statistical analyses:

All statistical analyses were performed using R. P-values were calculated using two-tailed tests, and Bonferroni corrections were conducted for multiple testing. The significance threshold was set to an adjusted P-value < 0.05. The effect of vessel caliber on the fluorescence intensity of *Myh11* was determined by one-way ANOVA, and the statistical significance between groups was tested with post-hoc tests (Turkey's test).

#### Supplemental Figures and Figure Legends:

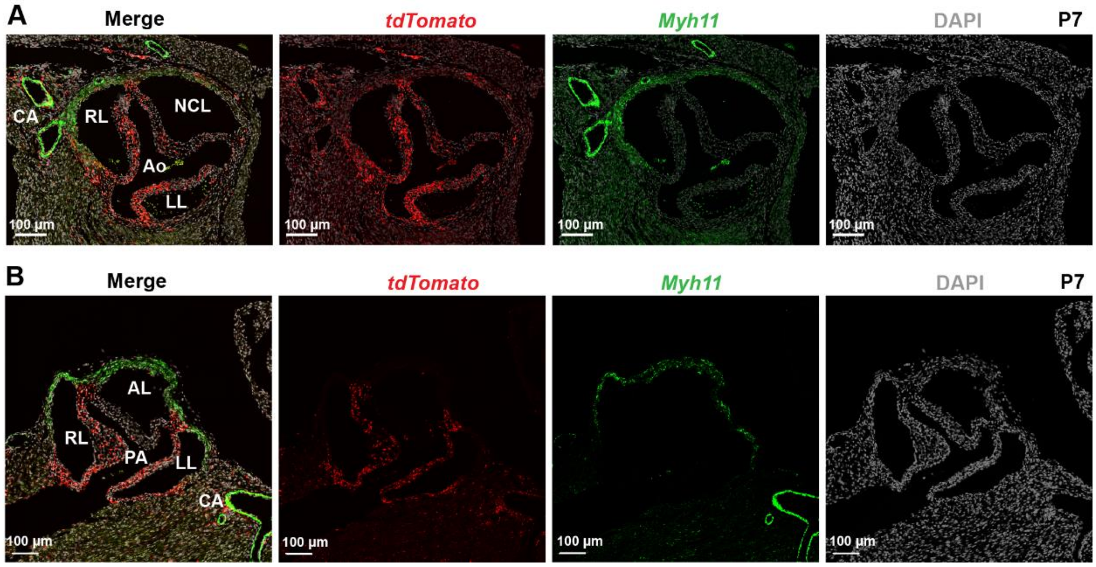


**Online Figure I. The distribution of CNCC-derived cells in the aortic and pulmonary valve leaflets. (A)** The distribution of CNCC-derived cells in the aortic valve leaflets. **(B)** The distribution of CNCC-derived cells in the pulmonary valve leaflets. Ao, aorta; AL, anterior leaflet; CA, coronary artery; Green, *Myh11*; Gray, DAPI; LL, left leaflet; NCL, noncoronary leaflet; PA, pulmonary artery; Red, *tdTomato*; RL, right leaflet

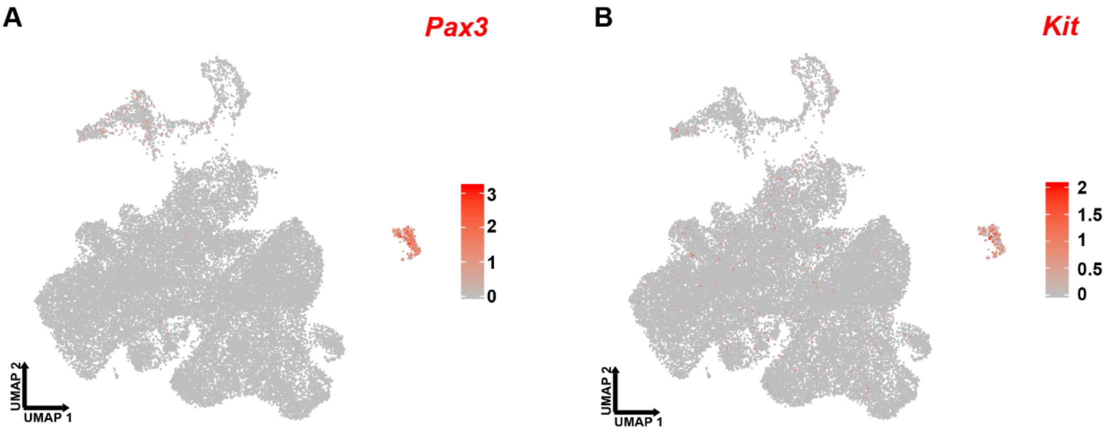


**Online Figure II. Melanocytes show high expression of the neural crest markers *Pax3* and *Kit*.**

#### Supplemental Table Legends:

**Online Table I. Summary of the sequencing metrics for the eight samples.**

**Online Table II. Molecular signatures of each cellular cluster.**

**Online Table III. Genes that were significantly regulated during the transition from pericytes to mVSMCs.**

**Online Table IV. Enriched Gene Ontology terms for the clusters of genes showing changes as a function of pseudotime.**

**Online Table V. UMI counts of marker genes for neuron and Schwann cells in the E10.5 sample.**

#### Legends for Supplemental Data:

**Online Data I. HTML page for interactive visualization of single-cell RNA-seq data in three-dimensional UMAP space.**
